# A comprehensive framework to capture the arcana of neuroimaging analysis

**DOI:** 10.1101/447649

**Authors:** Thomas G. Close, Phillip G. D. Ward, Francesco Sforazzini, Wojtek Goscinski, Zhaolin Chen, Gary F. Egan

## Abstract

Mastering the “arcana of neuroimaging analysis”, the obscure knowledge required to apply an appropriate combination of software tools and parameters to analyse a given neuroimaging dataset, is a time consuming process. Therefore, it is not typically feasible to invest the additional effort required generalise workflow implementations to accommodate for the various acquisition parameters, data storage conventions and computing environments in use at different research sites, limiting the reusability of published workflows.

We present a novel software framework, *Abstraction of Repository-Centric ANAlysis (Arcana)*, which enables the development of complex, “end-to-end” workflows that are adaptable to new analyses and portable to a wide range of computing infrastructures. Analysis templates for specific image types (e.g. MRI contrast) are implemented as Python classes, which define a range of potential derivatives and analysis methods. Arcana retrieves data from imaging repositories, which can be BIDS datasets, XNAT instances or plain directories, and stores selected derivatives and associated provenance back into a repository for reuse by subsequent analyses. Workflows are constructed using Nipype and can be executed on local workstations or in high performance computing environments. Generic analysis methods can be consolidated within common base classes to facilitate code-reuse and collaborative development, which can be specialised for study-specific requirements via class inheritance. Arcana provides a framework in which to develop unified neuroimaging workflows that can be reused across a wide range of research studies and sites.

## Introduction

Despite the availability of well-established neuroimaging analysis packages (Cox, 1996; Smith et al., 2004; Friston, 2007; Tournier et al., 2012), the arcana of neuroimaging analysis is substantial due to the range of available tools, tuneable parameters, and imaging sequences involved (Cusack et al., 2015). The distribution of complete “end-to-end” workflows, from acquired data to publication results, is necessary for routine reproduction because of the effort required to accurately reimplement such analyses (Kennedy, 2018). It is also difficult to adapt existing workflows to new studies without detailed knowledge of their design. Therefore, flexible, portable and complete workflows are important to promote reproduction and code reuse in neuroimaging research.

A barrier to designing portable and complete workflows is the heterogeneity of data storage conventions (Marcus et al., 2007; Das et al., 2012; Gorgolewski et al., 2016). To address this, the emerging Brain Imaging Data Standard (BIDS) (Gorgolewski et al., 2016) provides a way to standardise the storage of neuroimaging data in file system directories. BIDS specifies strict file and directory naming conventions, which facilitate the design of portable *BIDS Apps* (`bids-apps.neuroimaging.io`). However, for research groups with sufficient informatics support, software-managed repositories (Marcus et al., 2007; Das et al., 2012) can provide additional features, such as flexible access-control and automated pipelines. For published workflows, the choice of repository should be transparent in order to maximise their audience.

While neuroimaging analyses are generally amenable to standardisation (Kennedy, 2018), minor modifications are often required to accommodate idiosyncrasies of the acquisition protocols in use at different sites (Esteban et al., 2018). Workflows may require conditional logic in construction or execution to be portable. Nipype is a flexible Python framework for neuroimaging analysis in which workflows are constructed programmatically in Python (Gorgolewski et al., 2011). Programmatic construction allows for rich control-flow logic that is not readily available in alternative workflow frameworks (Cusack et al., 2015; Achterberg et al., 2016; Amstutz et al., 2016), and has been used to implement workflows that are robust to differences in fMRI protocols across a large number of sites (Esteban et al., 2018).

The trend towards large multi-site and multi-contrast datasets collected over a number of years (Van Essen et al., 2012; Thompson et al., 2014; Sudlow et al., 2015) presents additional challenges to workflow design. Analysis packages are constantly being developed and improved, so the state-of-the-art workflow for a particular analysis can change over time. Therefore, it is challenging to ensure workflows are applied consistently over the course of long studies (Cusack et al., 2015).

While analysis workflows for different contrasts and modalities are typically implemented independently, they can share common processing steps (e.g. non-linear registration to standard space, surface parcellation) and their outputs may need to be integrated to produce publication results. For large scale studies, which are typically processed on the cloud or high-performance computing (HPC) clusters, rerunning common segments can lead to significant increases in computation time and project cost. In addition, duplication of processing segments increases time for manual quality control (QC), making the reuse of intermediate derivatives a practical requirement for some large studies (Schreiber et al., 2018).

To maximise the reusability of neuroimaging workflows and avoid frequent reimplementation of standard analyses, workflow implementations should be flexible, extensible and applicable to a wide range of storage systems. In addition, in order to promote routine reproduction of neuroimaging studies, published workflow implementations should include the complete procedure, from acquired data to publication results. However, ensuring workflow implementations are flexible, portable and complete adds a high degree of complexity and effort to the design process.

Our objective was to extract common elements of repository-centric workflow design into an abstract framework to make it practical to implement flexible, portable and complete workflows for a wide range of neuroimaging analyses. *Abstraction of Repository-Centric ANAlysis (Arcana)* (`arcana. readthedocs.io`) is a Python framework for designing complex workflows in which modular Nipype pipelines operate on data stored in repositories. Intermediate derivatives are derived on demand, checking against stored provenance for required updates. Analyses can be applied to XNAT, BIDS and plain-directory repositories, and using Nipype’s execution plugins, run on workstations or be submitted as batch jobs to HPC schedulers. Arcana’s architecture, with programmatic workflow construction—yet clear delineation between analysis design and application—facilitates the implementation of complex workflows that are portable and complete.

The utility of the Arcana framework is demonstrated by the implementation of analysis suites for T1, T2* and diffusion weighted MRI (DWI) data and the application of DWI tractogram (Tournier et al., 2012) and vein masks (Ward et al., 2017) workflows to data collected from a healthy subject.

## Methods

### Framework overview

The separation of analysis design and application in the Arcana framework follows the conceptual divide between classes and objects in Object-Oriented (OO) software design. *Study* classes encapsulate types of data, such as scans of a specific imaging contrast or modality, with the suite of analysis methods that can be performed on them. Study objects apply the analysis suite defined in the Study class to a specific dataset.

The set of input data, the derivatives that can be derived from it, and methods that construct pipelines to derive the derivatives, are linked together by the *data specification table* class attribute of the Study (Figure 1). Likewise, free parameters used in pipeline construction are defined in the class’ *parameter specification table*. Class inheritance can be used to specialise analysis suites by overriding entries in the specification tables or pipeline constructor methods. Analysis suites for multi-modal data can be implemented by combining Study classes within *MultiStudy* classes.

**Fig. 1:**
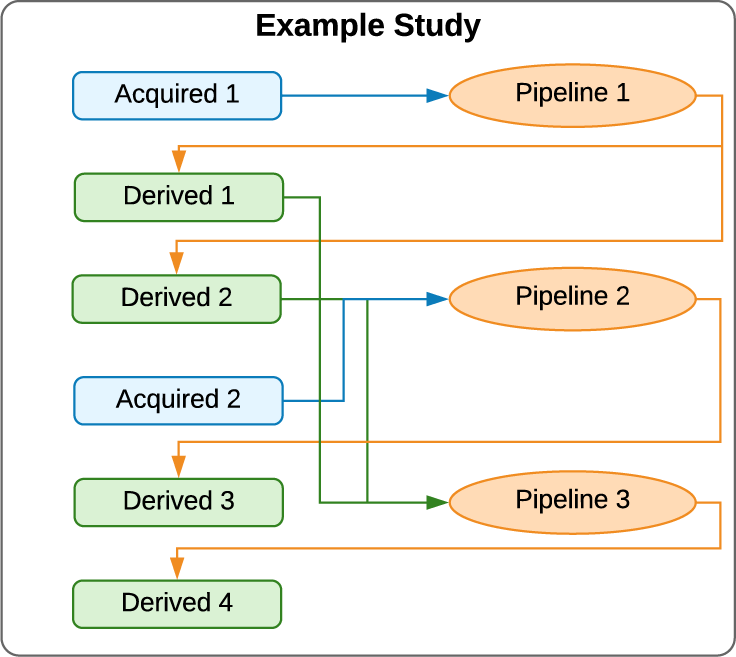
Example study. Blue boxes represent input data (filesets or fields) stored in a repository and green derivatives from that data stored alongside the original data. Orange ovals are pipelines that operate on data in the repository to derive the derivatives. Arrows represent data flows, i.e., inputs and outputs to pipelines

Analysis methods defined by a Study class are applied to a specific dataset by instantiating an object of the class and requesting a derivative listed in the class’ data specification table. At initialisation, a Study object is passed references to a *Repository*, a *Processor*, and an *Environment* objects, which define where and how data is stored and processed. When a derivative is requested, a Study object queries the Repository for intermediate derivatives that can be reused before constructing a workflow to produce the requested derivative. The manner in which the workflows are executed (i.e. single/multiprocess or via SLURM scheduler) is specified by the Processor and software modules required by the analysis are loaded by the Environment. Selected workflow products are stored back in the repository for reuse by subsequent analyses (Figure 2). Input data to a study are selected from repositories using criteria defined in *Input* objects passed to the Study object at initialisation and matched against entries in the class’ data specification table.

**Fig. 2:**
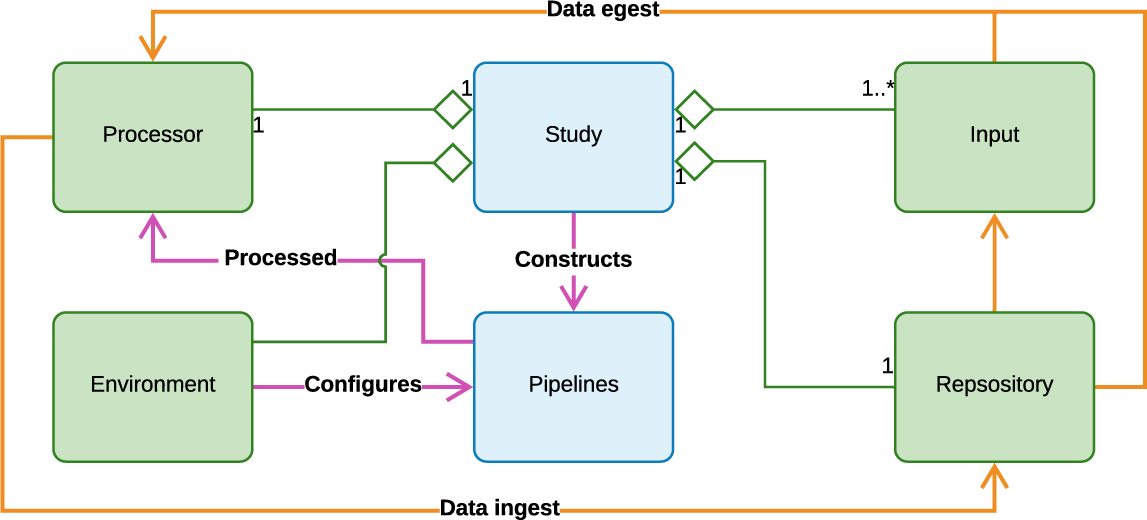
Unified Modelling Language (UML) diagram of information flow in the Arcana framework. Boxes: Python classes, blue=analysis-design, green=analysis-application. Arrows: orange=data, magenta=workflow description, diamond=aggregated-in. Study classes construct analysis pipelines, which are sent to the *Processor* to be processed. Input data is selected by *Input* objects and pulled to the compute environment to be processed along with existing intermediate derivatives. After the derivatives are pushed back to the repository.

### Analysis design: Study classes

Study classes encapsulate a study dataset (i.e. data collected across multiple subjects using the same acquisition protocol) with the suite of analytical methods that can be applied to the dataset. The hierarchy of a study dataset is assumed to have two levels, *subjects* and *sessions*, with each session for each subject corresponding to a specific *visit*, e.g. timepoint in longitudinal study. Derivatives can be created at any point in this hierarchy: per-session, per-subject, per-visit and per-study. Iteration over subjects and visits is handled implicitly by the framework. All Study classes must inherit from the *Study* base class and be created by the *StudyMetaClass* metaclass or subclasses thereof.

### Data and parameter specification tables

At the heart of each Study class is the data specification table, which is stored in the _*data_specs* class attribute and specifies the input and output data of the analysis, and all stages in between. There is a one-to-one relationship between entries in the data specification table and data that are stored in the repository, or will be stored if and when they are derived. Which intermediate derivatives to include in the data specification table, and therefore store in the repository, is left to the discretion of the researcher designing the analysis. However, as a general rule, derivatives that require manual QC or are likely to be reused between different branches of analysis should be included in the table.

Data specified in the data specification table can be of either a *fileset* or *field* type. Filesets represent single files, or sets of related files typically contained within a directory (e.g. a multi-volume DICOM dataset). Fields represent integer, floating point, datetime, or character string values. By default, a Field represents a single value but if the *array* flag is set, a Field represents a list of values. Each Fileset references a *FileFormat* object, which specifies the formats of the files in the set. File formats are explicitly registered by the researcher at design time using the *FileFormat.register(format)* class method to avoid conflicts where the same extension is used for different formats in different contexts.

Fileset and field specifications are passed to the data specification of a Study class via the *add_data_specs* class attribute as a list of named *FilesetSpec* and *FieldSpec* objects (Figure 3). Specifications for acquired data (i.e. input data to the study) are distinguished from derived data by using the *FilesetInputSpec* and *FieldInputSpec* subtypes. However, the distinction is fluid, with derived specifications able to be overridden by acquired specifications in subclasses or MultiStudy classes, and vice-versa, or passed inputs when the class is instantiated.

**Fig. 3:**
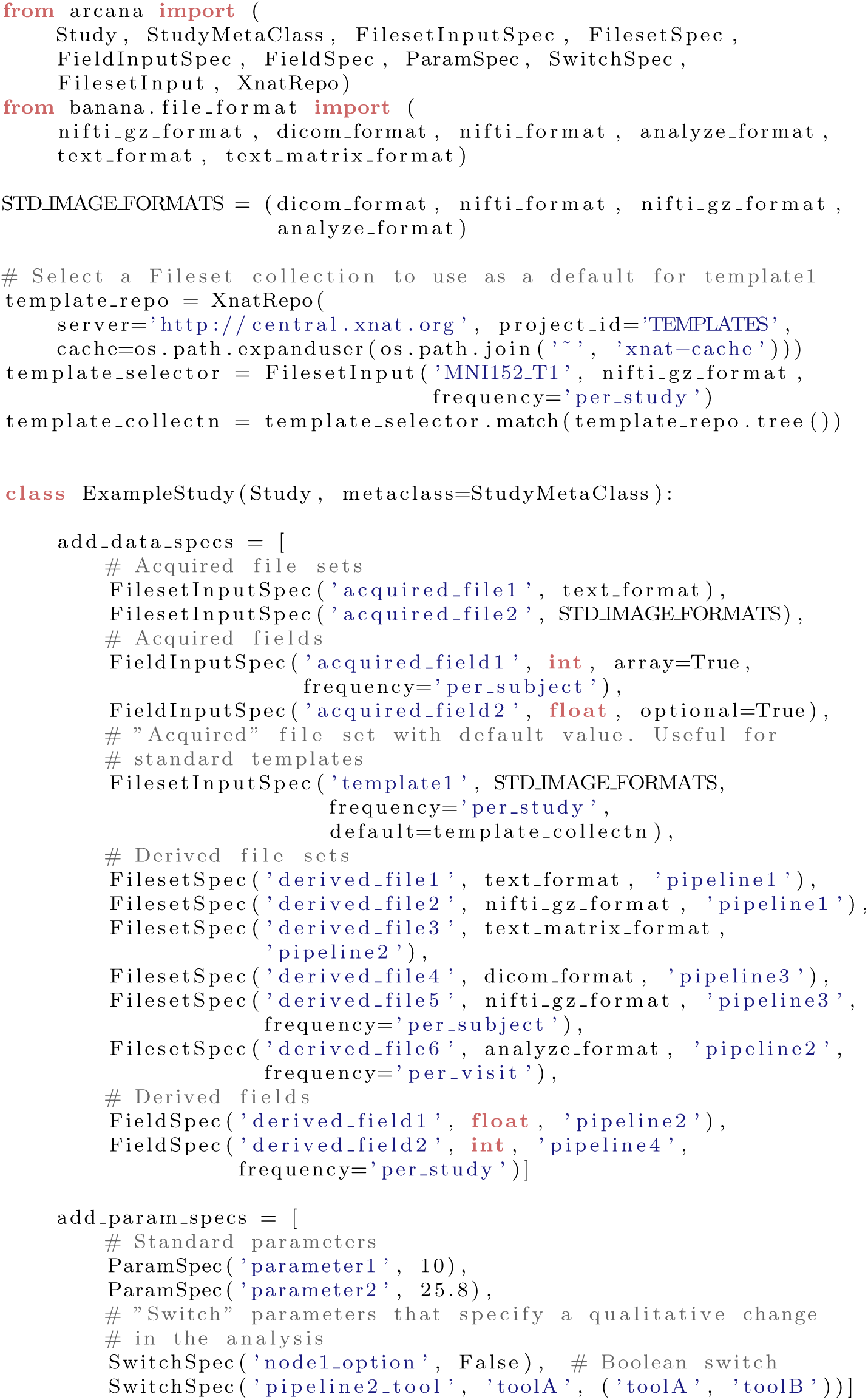
Example data and parameter specifications. The data specification specifies two input file sets, ‘one’ and ‘ten’ and ten derived file sets that can be derived from them, at least indirectly. Each derived data spec, specifies the name of the pipeline constructor that creates the pipeline that derives them. Parameter specifications specify a name and default value for free parameters of the Study class

All data specifications have a *frequency* attribute which specifies where the data sits in the hierarchy of the dataset and can take the values ’per_session’, ’per_subject, ’per_visit’ or ’per study’. In addition, derived specifications are passed the name of a method in the class that constructs the pipeline to derive them. Therefore, while a pipeline can have multiple outputs, each derivative is derived by only one pipeline.

For Study classes that correspond to a known type in the BIDS standard, a dictionary mapping their data specifications to default *BidsInput* or *BidsAssocInput* selectors can be specified in the *default*_*bids*_*inputs* class attribute. BidsInput specifies a primary scan in the BIDS standard using its *type, modality, format* and optionally the *task* it belongs to. BidsAssocInput is used to select associated files, such as field maps and diffusion encoding matrices relative to a primary scan. Default BIDS selectors are typically not provided with a value for the *task* keyword, which can be set at instantiation of the Study class with the *bids*_*task* keyword argument to enable the design of Study classes that are applicable to any BIDS task.

Similar to data specifications, parameter specifications are included in the Study class by providing a list of *ParamSpec* objects to the *add_param_specs* class attribute. ParamSpec objects are initialised with a name and default value. Special parameters that specify a qualitative change in the analysis, for example using ANTs registration (Avants et al., 2011) instead of FSL registration (Smith et al., 2004), are specified by the *SwitchSpec* subtype. SwitchSpecs take a name, default value and a list of accepted values at initialisation.

### Workflow design: pipeline constructor methods

Workflows are implemented in Arcana as a series of modular pipelines, which each perform a unit of the analysis (e.g. registration, brain extraction, quantitative susceptibility mapping). Pipelines are represented by *Pipeline* objects, which are thin wrappers around Nipype workflows to handle input and output connections, and namespace management. Each pipeline consists of a (typically small) graph of Nipype nodes, with each node wrapping a stand alone tool (e.g. FSL’s FLIRT) or analysis package function (e.g. SPM’s *coreg* tool). Pipelines source their inputs from, and sink their outputs to, entries in the data specification table, thereby linking the table together.

Pipelines are constructed dynamically by “pipeline constructor” methods of the study. Pipeline constructor methods are referenced by name in the *pipeline*_*name* argument of the FilesetSpec and FieldSpec objects of data specifications they derive. Pipeline constructor methods should only receive wild-card keyword arguments (e.g. my_pipeline(self, **name_maps)), and these arguments should be passed as a dictionary directly to the *name*_*maps* argument of the pipeline initialisation to enable inputs and outputs of the pipeline to be rerouted to alternative data specifications in modified constructor methods, typically in subclasses and multi-studies (see *Extension and specialisation by class inheritance* and *Implementing multi-modal studies*).

The syntax for pipeline construction is inspired by changes proposed for Nipype v2.0 (github.com/nipy/nipype/issues/2539). Within a pipeline constructor method, a Pipeline object is constructed by the *Study.pipeline* method (Figure 4). At initialisation, each pipeline is given a name, which must be unique amongst the pipelines constructed by the Study. The methods implemented by the pipeline can be characterised by providing a the list of citations and a text description to the *citations* and *desc* keyword arguments, respectively.

**Fig. 4:**
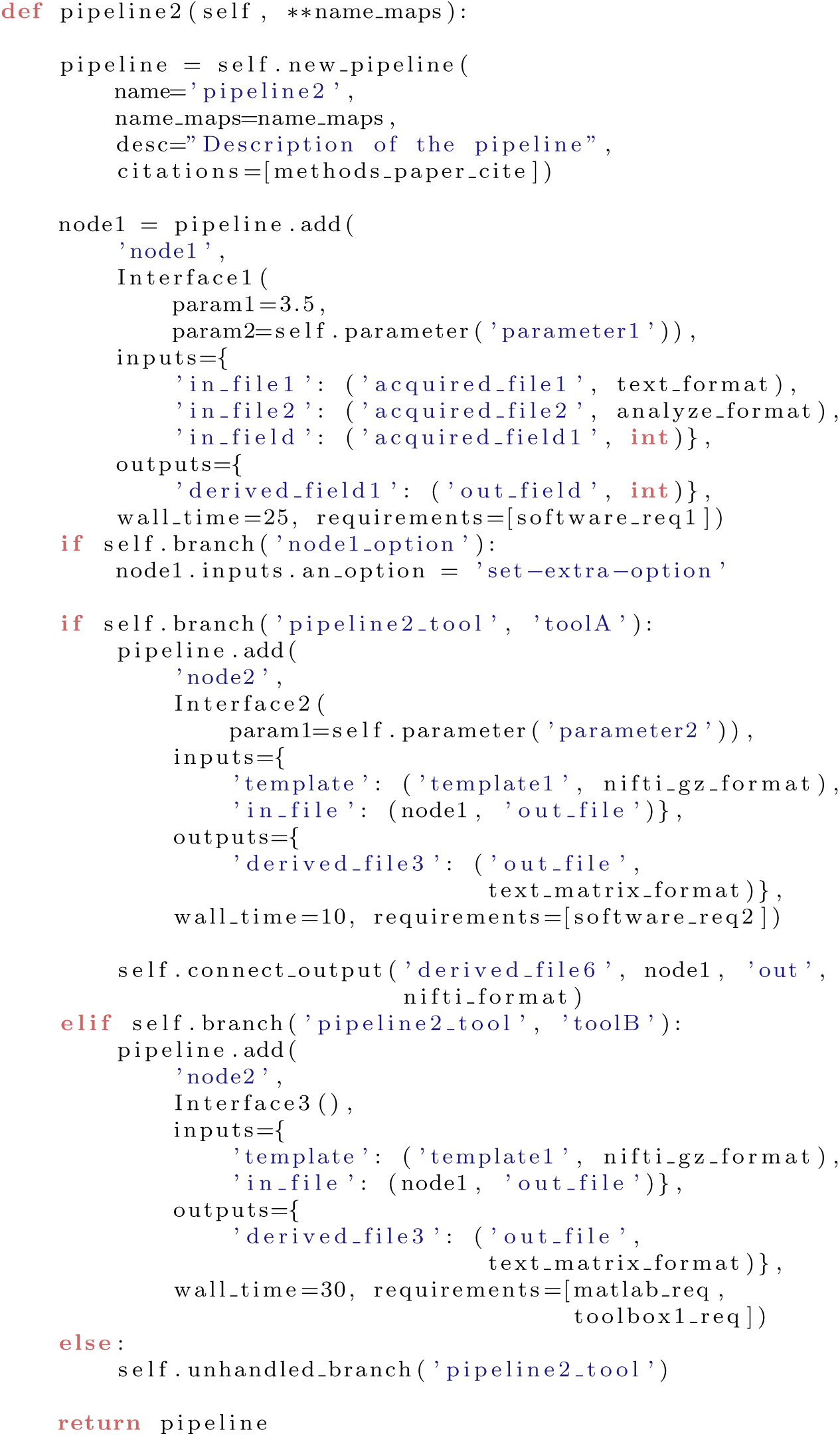
Example pipeline constructor method. Pipelines are created using the *pipeline* method of the Study class. Pipeline objects are thin wrappers around Nipype Workflow objects to in order manage the namespaces of the workflow’s inputs, outputs and nodes. Every pipeline constructor method should allow wildcard keyword arguments, which are passed to the *name*_*maps* argument of the pipeline initialisation. This allows pipeline constructors in sub and multi classes to map the inputs and outputs of the pipeline onto different data specifications.

The *add(name, interface)* method is used to add a node to a pipeline and takes a unique name for the node (within the pipeline) and a Nipype Interface object, and returns a reference to the newly added node. For clarity, is recommended to put all static inputs (i.e. parameters) of the interface as keyword arguments of the interface constructor (Figure 4). However, if an input conditionally depends on a parameter of the study it can be set via the *inputs* attribute of the node (e.g. my_node.inputs.my param = 1.0).

Node interfaces are connected to each other, and to inputs and outputs of the pipeline, by providing *inputs* and *outputs* keyword arguments to the *add* method. Both arguments take a dictionary. The keys of the *inputs* dictionary correspond to trait names in the node’s input interface, whereas the keys of the *outputs* dictionary correspond to names of entries in the data specification table. The values of both dictionaries are 2-tuples. For pipeline inputs, values of the *inputs* dictionary consist of a name of an entry in the data specification table and the format the input is expected in (i.e. a FileFormat for Fileset specifications or Python datatype for Field specifications). For pipeline outputs, values of the *outputs* dictionary consist of a trait name in the node’s output interface and the format the output data is generated in. For input connections from other nodes, values of the *inputs* dictionary consist of a reference to the upstream node and the name of a trait in the upstream node’s output interface (output connections to other nodes are implied by input connections of other nodes).

If the expected format of a pipeline input or generated format of a pipeline output does not match that of the corresponding study input or data specification, then a conversion node is implicitly connected to the pipeline by the framework to perform the required conversion. Pipeline inputs and outputs that are conditional on parameters or inputs provided to the study can be specified outside of the *add* method using the *connect*_*input* or *connect*_*output* methods, respectively. Conditional connections between nodes can be specified via the *connect* method of the pipeline (which simply calls the method of the same name in the underlying Nipype workflow).

Any external software packages required by a node should be referenced in the *requirements* keyword argument as a list of *Requirement* objects when the node added to the pipeline. Similarly, the expected memory requirements in MB and wall time for the node execution should be provided to the keyword arguments *memory*, and *wall*_*time*.

Iteration over subjects and visits is handled implicitly by Arcana and depends on the frequency of the pipeline’s inputs and outputs. To create a summary derivative (i.e. frequency != ’per_session’) from more frequent data, *Study.SUBJECT*_*ID* or *Study.VISIT*_*ID* should be passed to the *joinsource* keyword of the *add* method to join over subjects or visits, respectively. In this case, a JoinNode will be created instead of standard Node, which should be passed the additional keyword argument *joinfield* to specify the list of input traits to convert into lists to receive the joined input. Similarly, if the name of an input trait is provided (or list thereof) to the the *iterfield* keyword argument, then a MapNode will be created and the interface will be applied to all items of the list connected to that input (Gorgolewski et al., 2011). Additionally, the values of the subject and visit IDs are directly accessible as input fields of the pipeline named *Study.SUBJECT*_*ID* and *Study.VISIT*_*ID*, respectively.

Study parameters can be accessed during pipeline construction with the *Study.parameter(name)* method. If conditional logic is included in the workflow construction that alters the pipeline inputs, outputs or parameters then it should be controlled by a switch instead of a parameter. The analysis branch designated by a switch value should be tested with *Study.branch(name)* in the case of boolean switches and *Study.branch(name, ref*_*value)* in the case of string switches.

### Extension and specialisation by class inheritance

Because Arcana analyses are implemented as Python classes, class inheritance can be used to specialise existing analyses.

Instead of being set directly, the data and parameter specifications are set by the metaclass of the Study (i.e. StudyMetaClass) in order to combine them with corresponding specifications in the class’ bases. The combined data and parameter specifications are constructed by visiting the class’ bases in reverse method resolution order (MRO) and adding specifications from their *add*_*data*_*specs, add*_*param*_*specs* attributes, overriding previously added specifications with matching names. Note that in this scheme, specifications can only be appended or overridden but not removed by Study subclasses so as not to break workflows inherited from base classes.

Pipeline constructor methods can be overridden in subclasses like any Python method. Often the overriding method will call the superclass method to construct a pipeline, apply modifications and return the modified pipeline. In this scenario, references to the data specification in the superclass method can be mapped onto different entries in the table of the subclass by providing the *input*_*map* and *outpu_map* keyword arguments when calling the superclass method. Additionally the name of the pipeline can be altered by providing the *name* argument, which is useful when creating multiple pipeline constructor methods from a single base method. In these scenarios it is important to also pass the the wildcard keyword arguments of the overriding method to the *name_maps* keyword argument of the superclass method to allow the overriding method to be overridden in turn.

### Implementing multi-modal studies: MultiStudy classes

While basic Study classes are typically associated with single image modality or contrast, the analysis suites implemented by them can be integrated into multi-modal analysis by aggregating multiple Study classes (sub-studies) in a *MultiStudy* class. Analysis suites are integrated by joining the data specification tables of the sub-studies of a MultiStudy class. Entries in specification tables of sub-studies are joined by mapping them to a common entry in the specification table of the MultiStudy (Figure 5). This enables derivatives from one sub-study (e.g. brain extracted T1-weighted anatomical) to be referenced by workflows of other sub-studies (e.g. anatomically constrained DWI tractography).

**Fig. 5:**
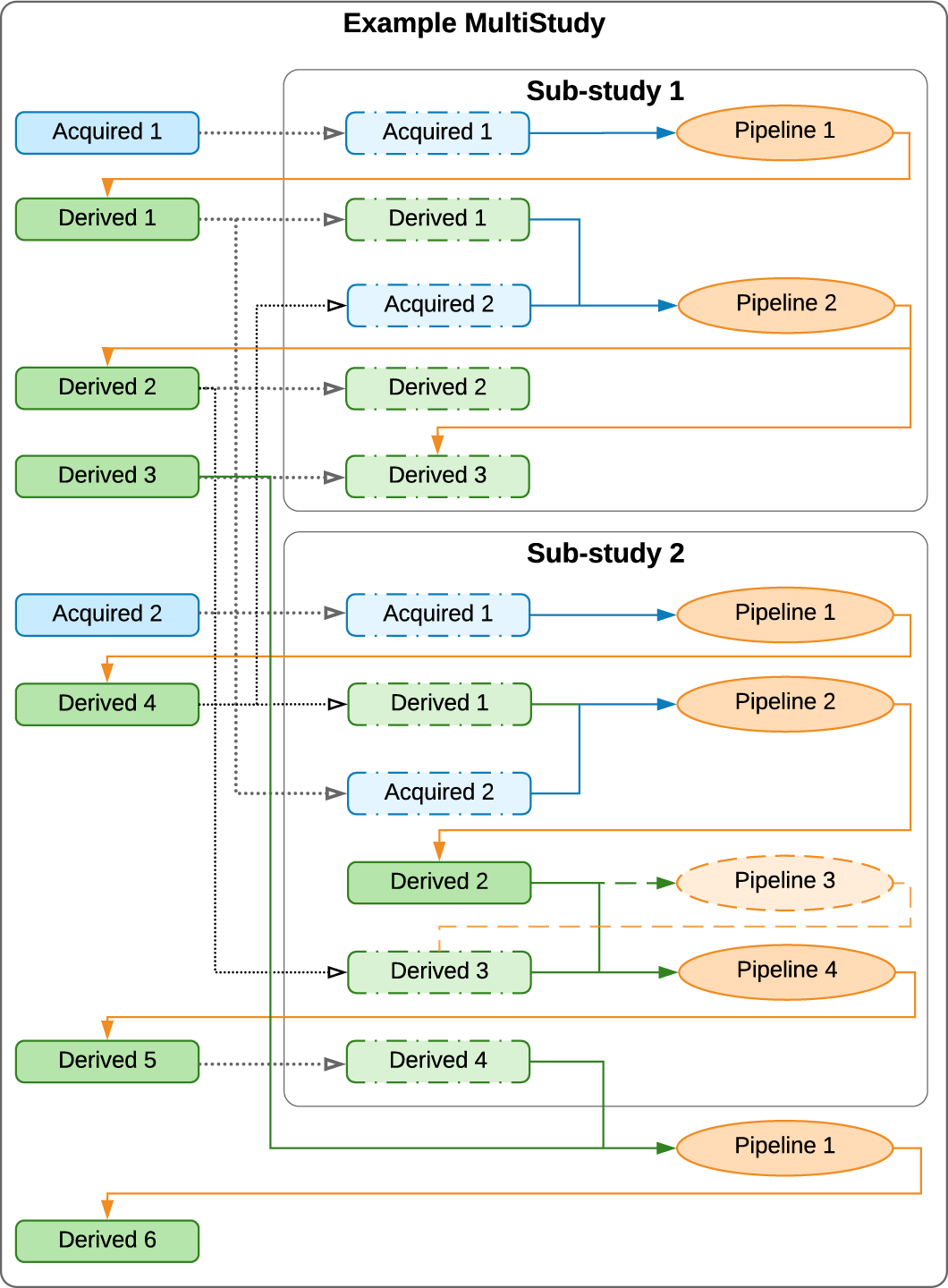
Example MultiStudy. Blue boxes represent input data (filesets or fields) and green derivatives. Orange ovals are pipelines. Blue and green arrows: pipeline inputs from study inputs and derived data, respectively. Orange arrows: outputs of pipelines. Dashed boxes represent data specifications in a sub-study that are present in the global namespace and mapped into the sub-study space, and dotted arrows the mappings. Sub-studies are linked by mapping the same data spec in the global space onto data specifications the multiple sub-study namespaces (e.g. *Derived 1, 2* and *3*). There are no restrictions between mapping study input and derivative data specifications: both input and derivative specifications can be mapped onto input data or derivative specifications in sub-studies. If a spec in the global spaced is mapped onto a derivative spec in the sub-study space, then the pipeline that generates that derivative in the sub-study will not run unless it generates other required derivatives (e.g. *Pipeline 3* in *Sub-Study 2*)

All MultiStudy classes must inherit from the Multi*Study* base class and be created by the Multi*StudyMetaClass* metaclass or subclasses thereof. As in the case of subclassing the standard Study class, additional data and parameter specifications can be added to the class via *add_data_specs* and *add_param_specs* respectively for additional analysis not included in the sub-studies.

Sub-studies are aggregated in the *sub-study specification table* of a Multi-Study class via a list of *SubStudySpec* objects in the *add*_*sub*_*study*_*specs* class attribute in the manner of data and parameter specifications. A SubStudySpec consists of a name, a Study class, and a *name map* dictionary. The name map dictionary maps data and parameter specification names from the sub-study namespace to the namespace of the MultiStudy class, i.e. the dictionary keys refer to entries in the sub-study specification table and the dictionary values refer to entries in the multi-study specification table.

Entries in the specification tables of sub-study classes that are not referenced in the sub study’s name map are implicitly mapped to the MultiStudy namespace by the MultiStudyMetaClass during construction of the MultiStudy class using the name of the sub-study as a prefix. If the implicitly mapped specification is derived, then its associated pipeline constructor method is also mapped into the MultiStudy namespace with a prefix. For example, *derived1* in *sub_study2* would be mapped to *sub_study2_derived1* along with the method *sub_study2_pipeline1*.

### Analysis application: Study instances

To apply the analysis blueprint specified in a Study class to a specific dataset, an instance of the Study class is created with details of where the data is stored (*Repository* module) the computing resources available to process it (*Processor* module) and the software installed in the environment it will be processed in (*Environment* module). A Study object controls the construction and execution of analysis workflows, and the flow of data to and from the repository (Figure 6).

**Fig. 6:**
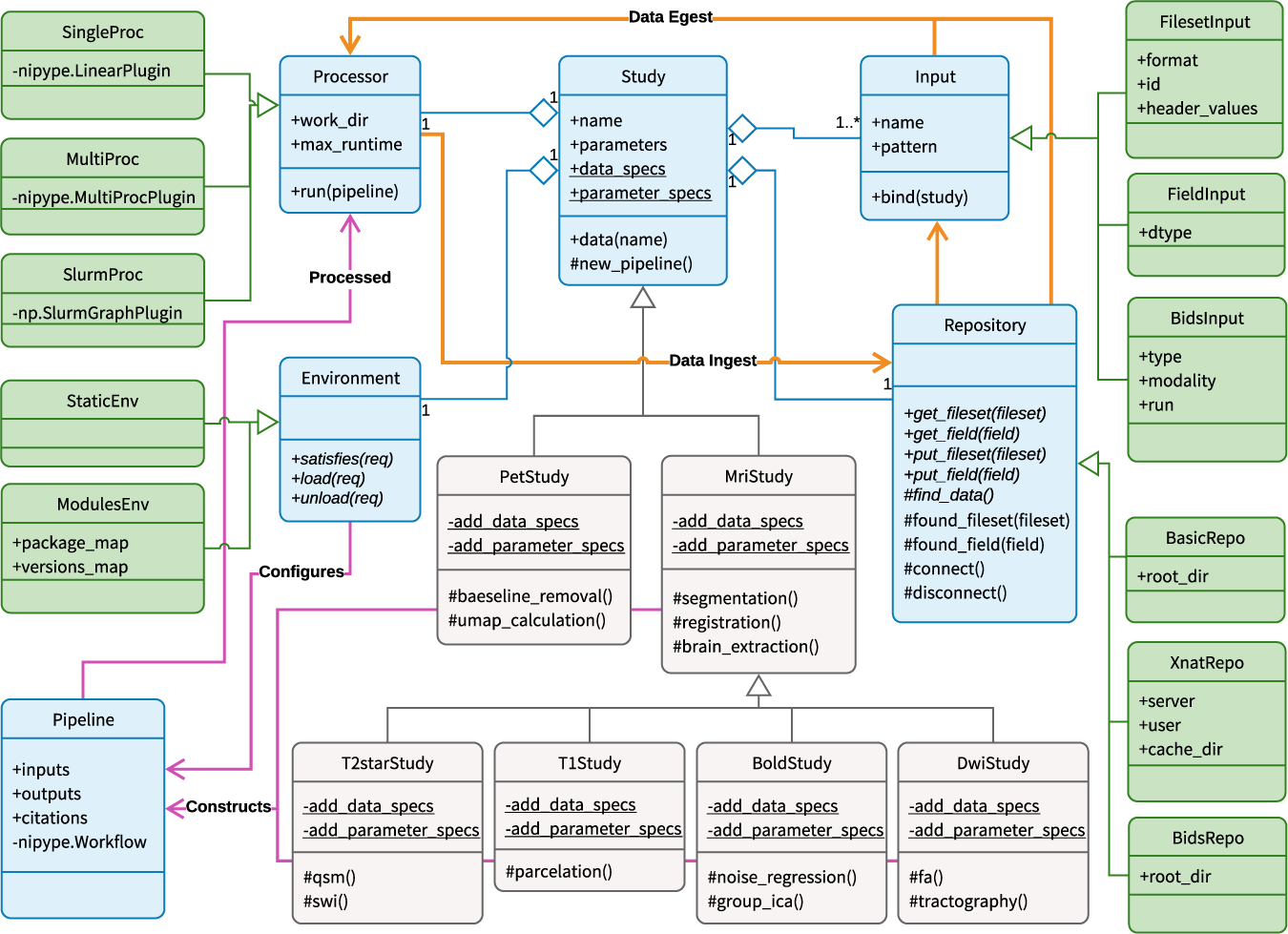
Detailed Unified Modelling Language (UML) diagram of information flow in the Arcana framework. Boxes: Python classes (blue=core, green=interchangeable modules, grey=example specialisations). Arrows: orange=data, magenta=workflow description, diamond=aggregated-in, triangle=subclass-of. Calling *data(name)* on a Study subclass constructs the requisite pipelines (as specified in *data_specs*) to produce the requested data, and sends them to the *Processor* to be processed. Data is selected by *Input* objects, pulled to the compute environment to be processed, and then the derivatives are pushed back to the repository. Repositories can be of plain directories, or BIDS or XNAT repositories

Each Study instance is assigned a name, which is used to differentiate its results from alternative analyses on the same dataset (e.g. with different parameterisations). Parameters are set on initialisation of the Study object along with the range of subject and visit IDs to be included in the analysis (if they are not provided then all IDs found in the repository are included). The remaining arguments passed to the Study object initialisation are the Repository, Processor and Environment modules to use and a list of *Input* objects to select input data from the repository and match it with the Study’s data specification table. (Figure 7).

**Fig. 7:**
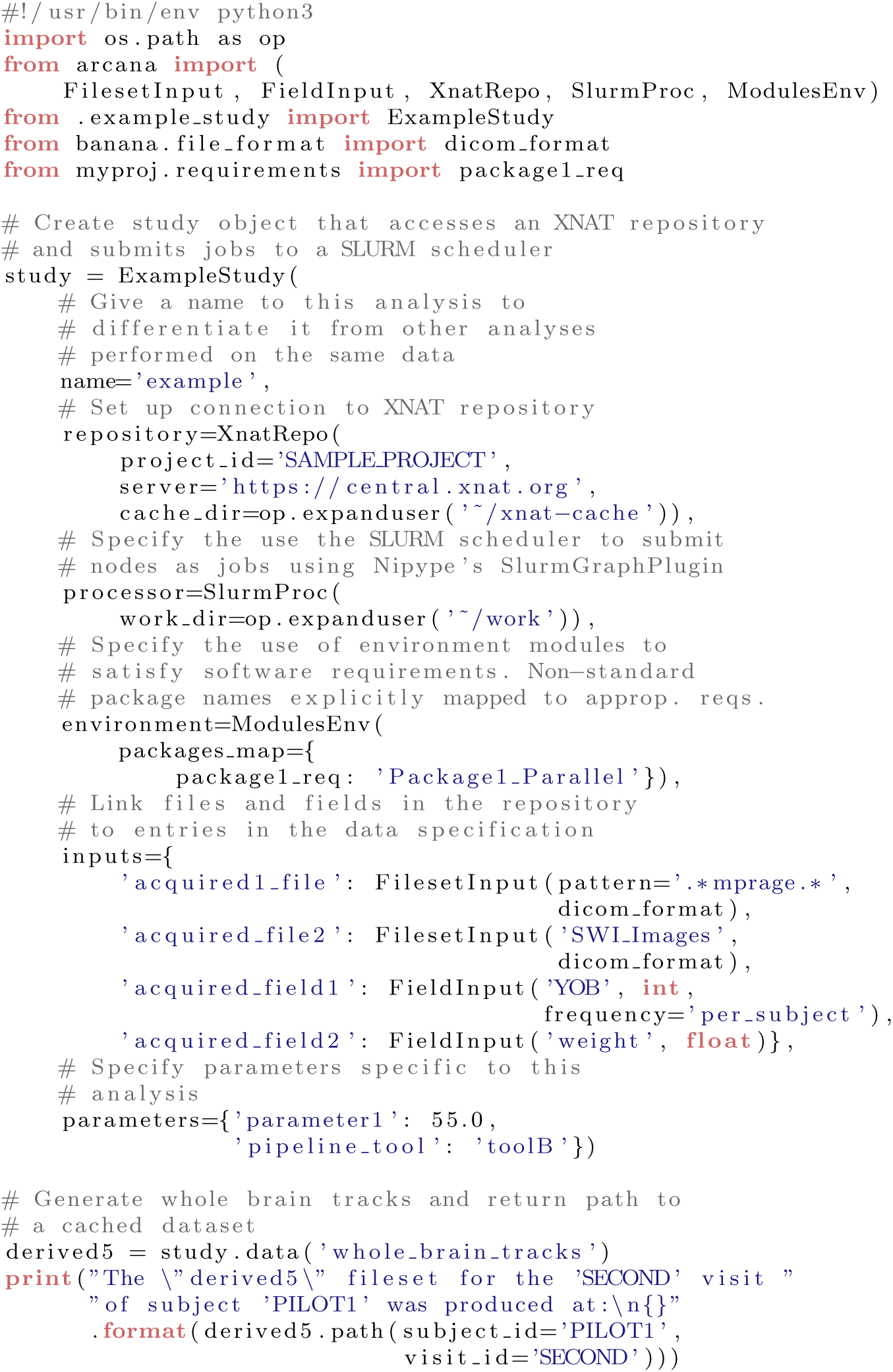
Example application of Study class to a dataset stored in an XNAT repository. Once the Study object has been initialised potential derivatives of the Study can be requested, and will be generated and stored in the repository if already present.

### Repository modules

In Arcana, repository access is encapsulated within modular *Repository* objects to enable switching between different repositories and repository types at analysis “application time” (Figure 2). There are currently three supported repository types, XNAT (Marcus et al., 2007), BIDS (Gorgolewski et al., 2016) and a “basic” format, which are encapsulated by *XnatRepo, BidsRepo*, and *BasicRepo* classes respectively.

In its most basic form, a “basic” repository, is just a file system directory containing the data to be processed for a single subject. For multi-subject studies, the root directory should contain separate subdirectories for each subject in the study, with subject IDs taken from the subdirectory names. If data was acquired for each subject over multiple visits, than an additional layer of nested subdirectories is included in the subject subdirectories, with the visit IDs taken from the subdirectory names. Note that the basic repository is similar to the BIDS format, however, there are no naming conventions in the basic repository, which enables prototyping and testing of analyses on loosely structured data.

Derivatives are stored by their specification name in a study-specific namespaces to avoid clashes with separate analyses. In basic repositories this namespace is a subdirectory named after the study nested within the lowest layer of the data tree. In BIDS repositories, the namespace is a subdirectory of the *derivatives* directory, again named after the study. In XNAT repositories, derivatives for each session are stored in separate *MrSession* objects alongside the primary session underneath its *Subject*, and are named <*primary-session-name*>_<*study-name*> (Table 1).

**Table 1:** Storage locations of derived data for each repository type. Derivatives are stored in separate namespaces for each Study instance to enable multiple analyses on the same datasets with different parameterisations. Where ‘…’ is the location of the directory or MrSession that holds the derivatives, *subj* = subject ID, *vis* = visit ID, *study* = name of the Study instance, and *pl* = name of pipeline.

| Datatype | Plain-directory | BIDS | XNAT |
| --- | --- | --- | --- |
| Derivatives | */subj/vis/study* | */derivatives/study/subj/vis* | */subj/vis_study* |
| Fields | *.../_fields_.json* | *.../_fields_.json* | MrSession XML |
| Provenance | *.../_prov_/_pl.json* | *.../_prov_/_pl.json* | *.../_prov_/_pl.json* |

Derived filesets are stored with the format specified in the study’s data specification. In plain-directory and BIDS repositories, fields are stored in a single JSON file named ‘fields .json’ in each derived session, and on XNAT they are stored in custom fields of the derived session. Provenance is stored in a ‘ prov ’ sub-directory (dataset on XNAT) of the derivatives directory (MrSession on XNAT) in separate JSON files for each pipeline named after the pipeline (Table 1).

Summary data (i.e. with *per subject, per visit*, and *per study* frequencies) are stored in specially named subjects and visits (e.g. ‘group’), the names for which are specified when the repository is initialised. For example, given in plain-directory repository using all as the summary name for both subjects and visits, per subject data for ‘subj1’ would be stored at <*root*>*/subj1/* group, per visit data for ‘visit1’ in <*root*>*/* group*/visit1*, and per study data in <*root*>*/* group/group (Table 1).

Each study can only have one repository in which derivatives are stored. However, a study can draw data from multiple auxiliary repositories, which are specified in the inputs passed to the study. When using multiple input repositories, subject and visit IDs will often need to be mapped from their values in the auxiliary repositories to the “ID space” of the study, which can be done by passing either by a dictionary or callable object (e.g. function) to the *subject*_*id*_*map* or *visit*_*id*_*map* keyword arguments during initialisation of a repository.

New repository modules for additional repository types can be implemented by extending the Repository abstract base class and implementing five abstract methods, *find*_*data, get*_*fileset, get*_*field, put*_*fileset* and *pu_field* (Table 2).

**Table 2:**
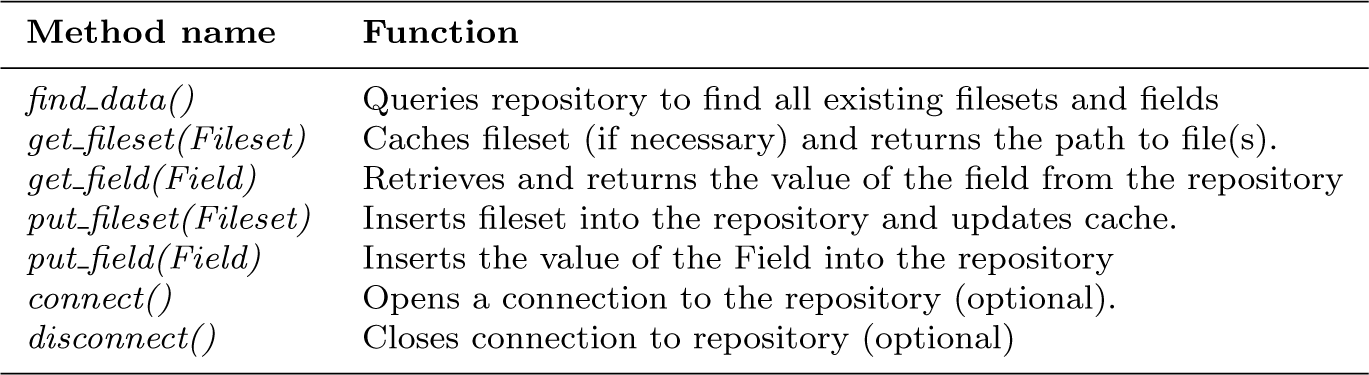
Abstract methods in the base Repository class that need to be implemented by platform-specific sub-classes.

### Study inputs

While derivatives generated by a Study object are named in accordance with the data specification of the Study class, arbitrary naming conventions can be used for input datasets and fields. A selection stage matches input data to entries in the data specification table of the Study class. The criteria for these selections are passed to the the *inputs* argument of the Study object at instantiation in a dictionary mapping data specification names to *FilesetInput* and *FieldInput* objects, and are required to match exactly one fileset or field in every session included in the study.

Matching is typically performed on file names (dataset labels for XNAT repositories) and field names. If the names are inconsistent across the study then regular expressions can be used instead of exact matches with the *pattern* keyword argument. Additional criteria can be used to distinguish cases where multiple filesets in the session match the pattern provided, such as DICOM header values, or the order and ID of the dataset.

Inputs can be drawn from auxiliary repositories by providing alternative Repository instances to the *repository* keyword of the Input. Care should be taken to ensure that the subject and visit ID schemes will map correctly to that of the primary repository (see *Repository modules*). If the primary repository is empty (i.e. all inputs come from auxiliary repositories) then explicit *subject*_*ids, visit ids* need to be provided and the *fill*_*tree* flag set when initialising the Study.

If the data to select has been derived from an alternative Arcana Study instance, then the name of the alternative study can be passed to the *from*_*study* keyword argument provided to the selector. There are no restrictions on selecting any data, derived or otherwise, to match input or derived specifications. For example, it is possible to deliberately skip analysis steps by selecting an output of an early step in a previous run as a match for a later derived spec although this is not recommended as standard practice.

Specific files and fields that are not stored within a repository can be passed as inputs in *FilesetCollection* and *FieldCollection* objects. Collection objects reference an entry in the data specification table and contain a single Fileset or Field for every session (or subject/visit/study depending on the frequency of the corresponding data specifcation). Collection objects can be used to pass reference atlases and templates as inputs to analyses. They can also be set as the default for a data specification via the *default* keyword argument. However, for the sake of portability, default inputs should be restricted to data in publically accessible repositories or those included in standard software packages (e.g. FSL).

When using BIDS repositories, the selection stage is typically already included in the data specification (see *Data and parameter specifications)* so inputs do not need to be provided to the initialisation of the Study. However, BidsInput and BidsAssocInput objects can be provided to override the default selections if required.

### Processor modules

Processor modules control how pipelines generated by a Study are executed. There currently three Processor modules implemented in Arcana: *SingleProc, MultiProc, SlurmProc*, which wrap the LinearPlugin, MultiProcPlugin and SlurmPlugin Nipype execution plugins, respectively. The main task performed by the processor, as separate from the Nipype execution plugin it wraps, is to determine which pipelines need to be run and link them into a single workflow. Since this logic is implemented in the Processor abstract base class, wrapping additional Nipype plugins as required is trivial.

A Processor is used internally by a Study instance to execute pipelines to derive derivatives requested from the data specification by the *data(name[, name,…])* method (Figure 2). The first step in this procedure is to query the repository tree for all data and provenance associated with the study. Sessions for which the requested outputs of the pipeline are already present in the repository, and the stored provenance matches the current parameters of the study, excluded from the list to process. For the remaining sessions to process, inputs of the pipeline that are derivatives themselves are added to the stack of requested derivatives. This procedure is repeated recursively until there are no sessions to process or all inputs to the pipeline are study inputs at a given depth.

When a pipeline is processed it is connected to source and sink nodes, which get and put the pipeline inputs and outputs from and to a repository, respectively. Separate source and sink nodes are used for each data frequency (i.e. per-session, per-subject, per-visit, per-study). If implicit file format conversion is required (i.e. the input or output format differs from the data specification) then additional format converter nodes are inserted after the source nodes or before the sink nodes. Iterator nodes that iterate over the required subjects and visits are connected to the sources, and “deiterator” nodes that join over subjects and visits are connected to the sink nodes. Final nodes of upstream pipelines are connected to downstream iterator nodes in order to create a single workflow, which is then executed using the Nipype execution plugin.

Provenance is stored for each pipeline run alongside the generated derivatives and consists of parameter values used by the pipeline, software versions used by the pipeline, interface parameters, a graph representation of the underlying Nipype workflow, checksums of inputs, version of Arcana used, version of Nipype used, and any subject and visit IDs that were joined over in the workflow.

For subsequent analyses, changes with respect to any of the stored provenance values will be flagged as a mismatch, with the exception of interface package versions (e.g. Nipype or Banana versions). How mismatches are handled depends on the *reprocess* flag passed to the Processor. If *reprocess* is true then the sessions with mismatching provenance will be reprocessed, otherwise if *reprocess* is false (the default) an exception will be raised. Changes with respect to any element in the provenance JSON document can be ignored by providing a list of JSONPath queries to the *prov ignore* keyword argument.

### Environment modules

The software packages installed on the system that are required by a study’s workflows (e.g. FSL, SPM) are detected and managed by the Environment object passed to the study at initialisation. There are currently two types of Environment class implemented in Arcana: *StaticEnv* and *ModulesEnv*.

*ModulesEnv* objects can be used if environment modules (Furlani, 1991) are installed on the system (typical on many HPC systems). In this case, environment modules are loaded before a workflow node is run, based on the requirements specified for the node during construction of the pipeline (see *Pipeline constructors*), and then unloaded afterwards. Mappings from non-standard module names installed on the system to those expected by the Study can be can be passed as a dictionary to the *packages*_*map* keyword argument at initialisation. Likewise non-standard versions for packages can be mapped onto the versioning system of a requirement using the *versions*_*map* keyword argument.

*StaticEnv* objects don’t actively manage the environment, and instead only check the current environment for appropriate software versions before running requested workflows.

### Acquisition of test dataset

A healthy volunteer was scanned using a 3T Siemens Skyra with a 32-channel head and neck coil to demonstrate the application of analyses implemented in Arcana. The protocol was a T1-weighted MPRAGE (1mm contiguous, matrix size 256×240×192, FOV 256×240×192, TE = 2.13ms, TR, = 2300ms, TI = 900ms, bandwidth = 230Hz/pixel), GRE (1.8 mm contiguous, matrix size 256×232×72, FOV 230×208×130, TE = 20ms, TR, = 30ms, bandwidth = 120Hz/pixel), and diffusion MRI (2 mm contiguous, matrix size 110×100× 60, FOV 256× 240× 192, TE = 95ms, TR, = 8200ms, 33 diffusion directions with b = 1500 mm2/s and 3 b=0, bandwidth = 781Hz/pixel).

## Results

The Arcana framework is distributed as a publicly available software package via GitHub (github.com/MonashBI/arcana) and the Python Package Index (PyPI) (pypi.org/project/arcana/). Study classes for T1, T2* and diffusion weighted MRI data have been implemented as part of the *Biomedical imging ANAlysis iN Arcana (Banana)* package (github.com/MonashBI/banana; pypi.org./project/banana). All three classes, *T1Study, T2starStudy* and *DwiStudy*, inherit generic image analysis methods, such as registration and brain extraction, from the base class *MriStudy*.

The DwiStudy class implements the extraction of diffusion tensor metrics, FA and ADC, as well as whole-brain tractography using streamlines tracking from the MRtrix toolbox (Tournier et al., 2010, 2012) (Figure 8).

**Fig. 8:**
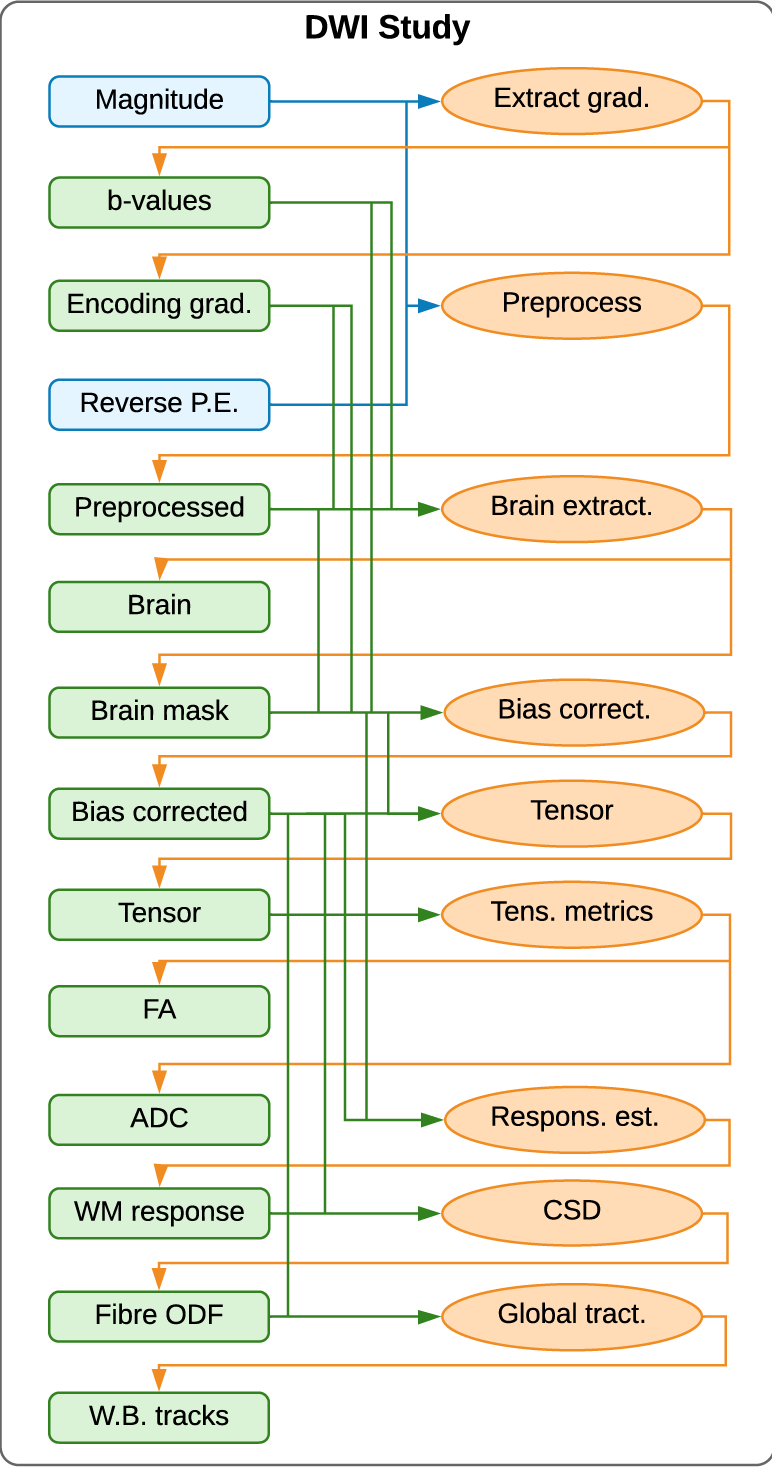
Example diffusion-weighted MRI (DWI) study, which can derive tensor metrics, fractional anisotropy (FA) and apparent diffusion coefficient (ADC) as well as streamlines fibre tracking. Blue boxes: acquired (input) data (filesets or fields). Green boxes: derivatives. Orange ovals: pipelines. Blue and green arrows: acquired and derived inputs to pipelines, respectively. Orange arrows: outputs of pipelines. The DWI magnitude image is preprocessed for motion correction and EPI distortions masked and bias corrected. From the bias corrected image two branches of analysis can be performed using the same intermediate derivatives: FA and ADC and/or streamlines fibre tracking.

The *T2starStudy* class implements an algorithm to generate *composite vein images* (Ward et al., 2018) and vein masks (Ward et al., 2017) from the combination of vein atlases derived from manual tracings with Quantitative Susceptibility Mapping (QSM) and Susceptible Weighted Imaging (SWI) images derived from the T2*-weighted acquisition (Figure 9).

**Fig. 9:**
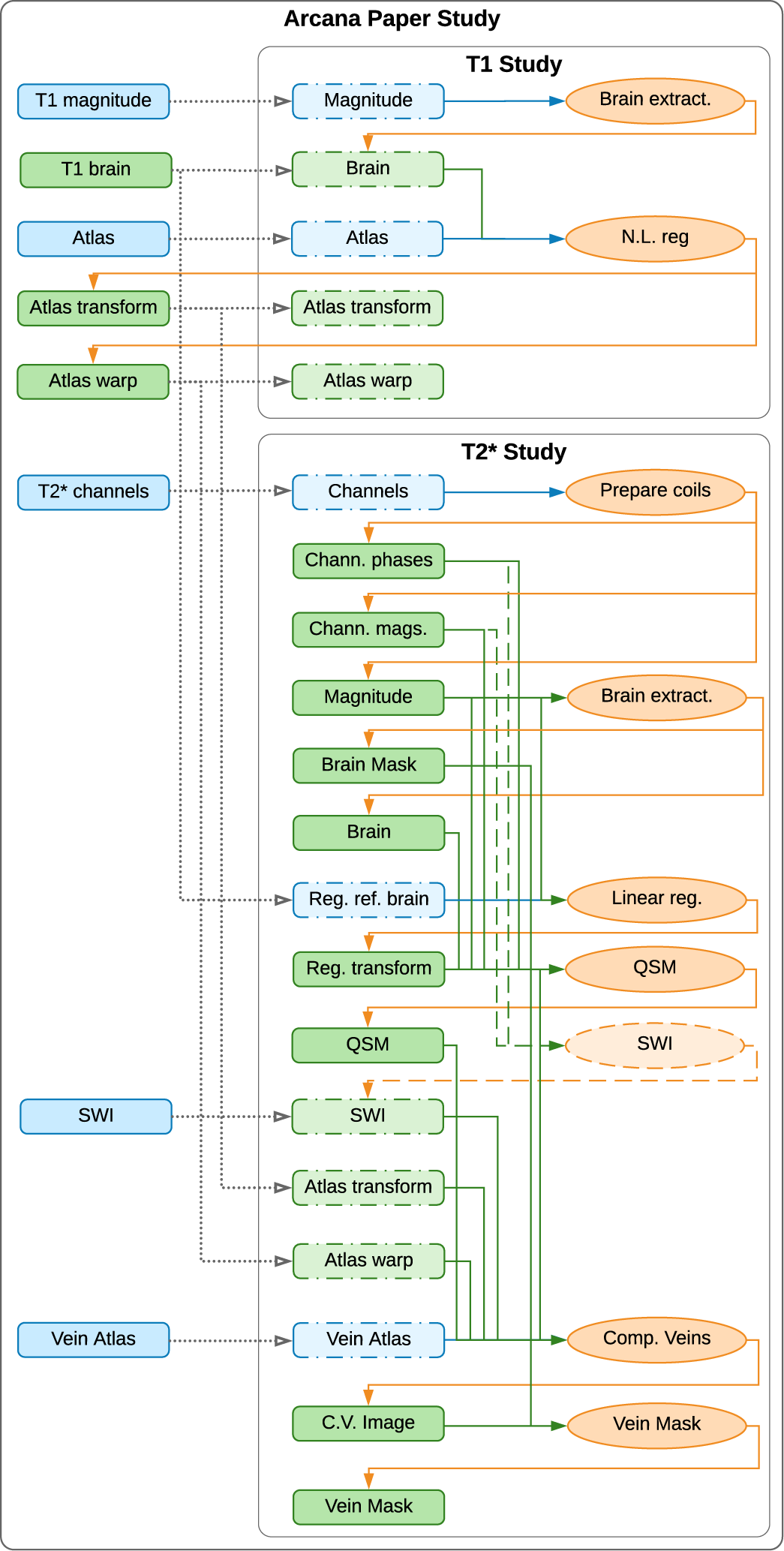
Combined T2*/T1-weighted studies within *ArcanaPaper* MultiStudy class, which can derive vein masks by combining Quantitative Susceptibility Mapping (QSM) and Susceptible Weighted Imaging (SWI) contrasts with a manual atlas. Blue boxes: input data (filesets or fields). Green boxes: derivatives. Orange ovals: pipelines. Blue and green arrows: inputs to pipelines from input and derived data, respectively. Orange arrows: outputs of pipelines. Dashed boxes represent data specifications in a sub-study that are present in the global namespace and mapped into the sub-study space, and dotted arrows the mappings. The acquired T1-weighted image is mapped to both the *magnitude* spec of the T1-weighted sub-study and the *registration reference* spec of the T2*-weighted sub-study. The nonlinear transformation from subject to atlas space are mapped from the T1-weighted sub-study and combined with the linear registration between T1-weighted and T2*-weighted images, QSM and SWI images are combined to produce the composite-vein image. In this instance, the SWI acquired from the scanner console is passed as an input to the derived *SWI* specification, overriding the *SWI* pipeline that would otherwise generate it (dashed oval).

The T1Study, T2starStudy and DwiStudy classes are aggregated into a single MultiStudy class, *ArcanaPaper* (Supplementary material), which is specialised to produce the figures in the Results section of this manuscript. In order to warp the vein atlases to the subject space for comparison with the SWI and QSM images, nonlinear registration to Montreal Neurological Institute templates (Grabner et al., 2006) is performed in the T1Study. The transform and warp field from this registration are then mapped onto the ‘coreg_to_atlas_mat‘ and ‘coreg_to_atla_warp’ specifications in the T2starStudy, as the registration of T2*-weighted images to the MNI template is typically poor. These transforms are combined with the linear transform from the brain-extracted T2*-weighted magnitude image to the brain extracted T1-weighted image, which requires the ‘brain’ specification in the T1Study to be mapped to the ‘coreg_ref_brain‘ specification in the T2starStudy.

The SWI image reconstructed on the scanner console is substituted for the *SWI* derivative produced by the *SWI* pipeline of the T2starStudy. For ease of comparison with the QSM and vein images produced by T2starStudy, this SWI image is brain extracted in a separate MriStudy sub-study of ArcanaPaper.

The *vein*_*fig, fa*_*adc*_*fig* and *tractography*_*fig* methods implemented in the ArcanaPaper class were applied to the test dataset to produce Figure 10, 11, and 12, respectively. Figure 10 displays composite vein images and vein masks for the healthy volunteer along with the SWI and QSM intermediate derivatives. The derived vein images are comparable to those generated by the original implementation (Ward et al., 2018).

**Fig. 10:**
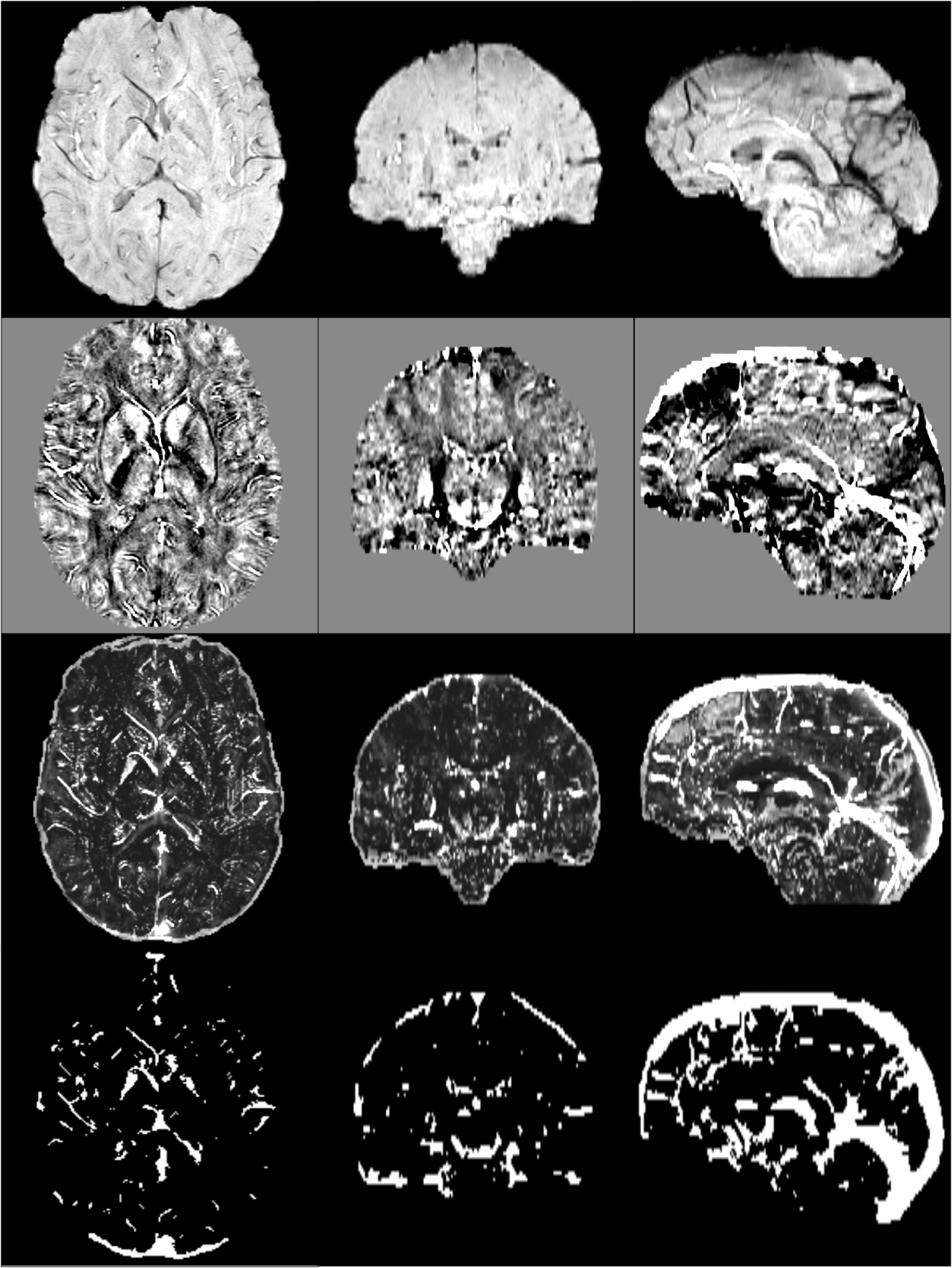
Composite vein image (*third row*) constructed by combining susceptibility weighted imaging (SWI) (*top row*), quantitative susceptibility mapping (QSM) (*second row*) and a vein atlas from manual tracings. A vein mask was then generated (*bottom row*) from the composite vein image. *left column*: axial slices. *centre column*: coronal slices. *right column*: sagittal slices.

**Fig. 11:**
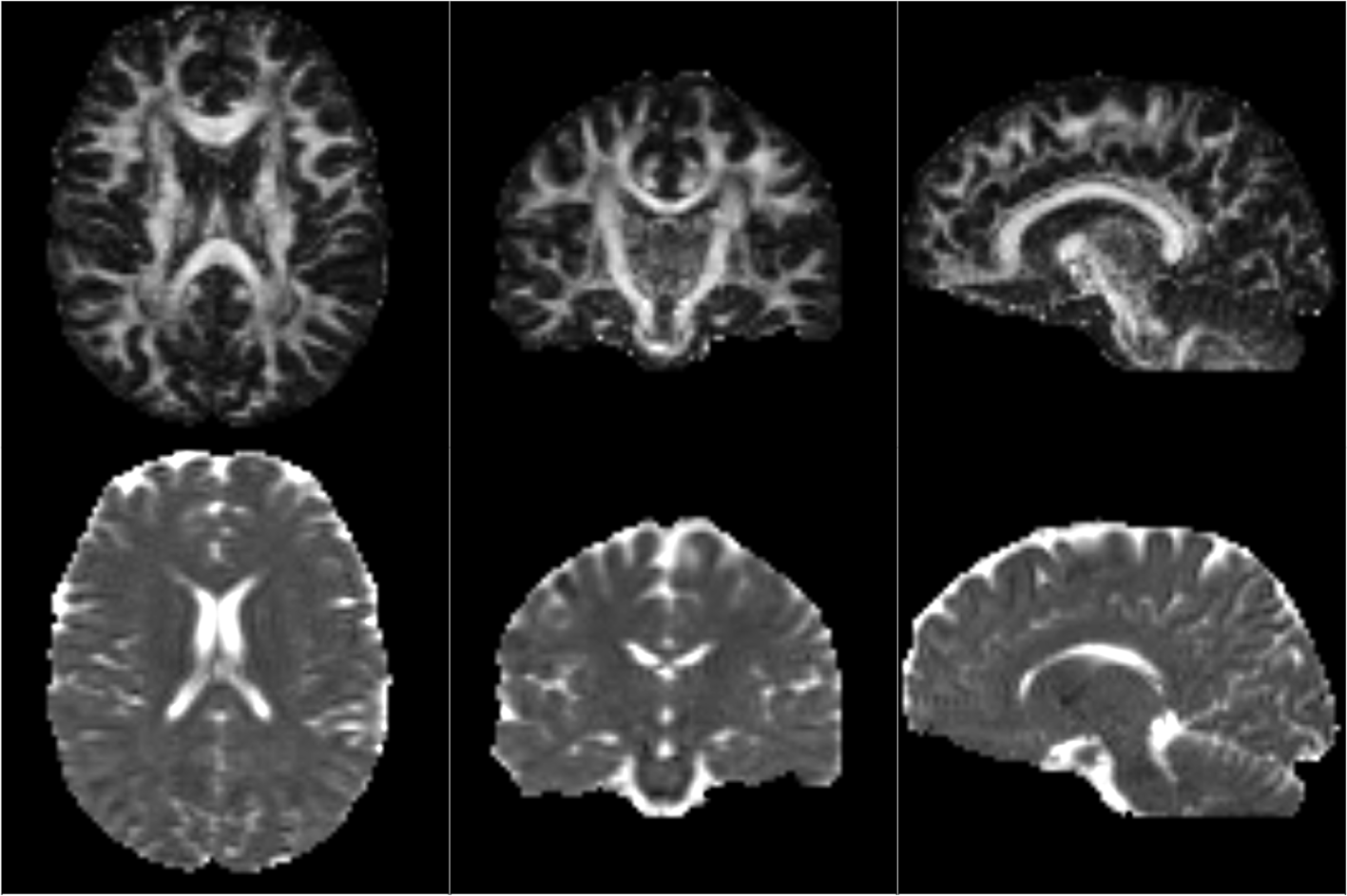
Fractional Anisotropy (FA) *(top row)* and Apparent Diffusion Coefficient (ADC) (*bottom rowI)* derived from diffusion MRI data. *Left column:* axial midline slices. *Middle column:* coronal midline slices. *Left column*: sagittal midline slices.

**Fig. 12:**
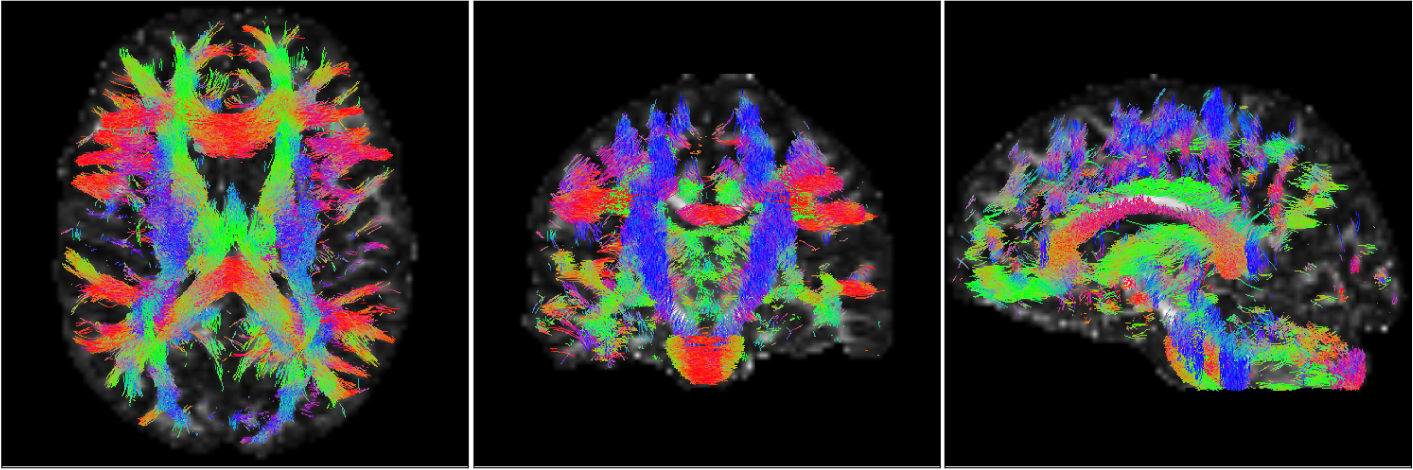
Global tractography performed using the MRtrix toolbox. Probabilistic streamlines generated with the iFOD2 algorithm from fibre Orientation Distribution Function (fODF) estimated from diffusion MRI datasets using Constrained Spherical Deconvolution (CSD). Streamlines are colour-encoded by orientation: green=anterior-posterior, blue=inferior-superior, red=left-right. *Left panel:* axial midline slice. *Middle panel:* coronal midline slice. *Left panel*: sagittal midline slice.

Figure 11 displays the FA and ADC maps derived from the diffusion MRI acquisition. The FA map shows high intensity in known white matter tracts and low intensities in known grey matter regions. The ADC map shows high intensity in cortical spinal fluid and low intensity through the rest of the brain. Figure 12 displays the global tractography derived from the DWI acquisition. The streamlines follow well known white matter tracts such as the corticospinal, fasciculus and corpus callosum. Intermediate derivatives derived for the FA and ADC analysis, including the preprocessed and bias-corrected DWI image and a whole brain mask, were reused in the generation of the streamlines.

## Discussion

We present Arcana, a software framework to facilitate the development of comprehensive analysis suites for neuroimaging data that implement complete workflows from repository data to publication results. The encapsulation of repository data and workflow generation in Arcana enables researchers to create robust workflows while focussing on the core logic of their analysis. Arcana’s modular pipeline and OO architecture promotes code reuse between different workflows by facilitating the sharing of common segments (e.g. registration, segmentation). The clear separation of analysis design from its application leads to portable workflows, which can be applied to datasets stored in a number of storage systems. In addition, the management of intermediate derivatives, provenance and software versioning, coupled with ability to submit jobs to HPC clusters, enables workflows implemented in Arcana to scale to large datasets. Arcana thereby enables researchers to quickly prototype analysis suites on local workstations that can be deployed on enterprise-scale infrastructure without modification.

Software frameworks (Yacoub and Ammar, 2004) have been successful in improving code quality and efficiency of development in a variety of contexts (Moore et al., 2008; White, 2012; Abadi et al., 2016). By factoring out common elements, only features that are specific to the given application need to be implemented by the analysis designer, and the common elements become battle hardened through repeated use. Arcana handles many of the menial tasks involved with workflow implementation, such as data retrieval and storage, format conversions, and provenance, reducing the time and effort required to implement robust workflows from acquired data to publication results.

An oft-repeated mantra in the open-source software movement dubbed Linus’ Law is that “given a large enough beta-tester and co-developer base, almost every problem will be characterized quickly and the fix obvious to someone” or more compactly, “given enough eyeballs, all bugs are shallow” (Raymond, 1999). Given the size of the neuroimaging research community, there are a large number of potential beta-testers and co-developers. However, it has been difficult for researchers to collaborate on the same code-base due to slight differences in acquisition protocols, storage conventions, researcher preferences, and study requirements.

The flexibility and portability of the Arcana framework increases the feasibility of community collaborations on workflow implementations. The improvement of code quality in larger community efforts, due to more eyeballs to detect and fix errors, has the potential to form a reinforcing cycle where more developers are attracted to the project. To these ends, the Banana code repository on GitHub (github.com/MonashBI/banana.git) is proposed as a code base for communal development of biomedical imaging workflows using Arcana.

A level of proficiency in Python OO design is required to design new analyses in Arcana, which may preclude inexperienced programmers. However, only a basic knowledge of Python is required to apply existing analyses to new datasets. Furthermore, a number of example Study classes have been implemented, which can guide the hand of analysis designers. Arcana imposes a consistent structure on workflows implemented within it, making the code easier to understand for developers who are familiar with the framework. In addition, class inheritance provides a manageable way to adapt and extend to existing analyses and highlights where modified analyses differ from standard procedures.

MR contrast-specific analyses are implemented in Banana via a chain of successively specialised Study sub-classes (e.g. MRI>EPI>DWI) to enable generic processing steps (e.g. registration) to be shared between classes. While not necessary, it is recommended to create a subclass specific to the research study in question and aggregate all related analysis within it, since such classes can be applied to alternate datasets in order to reproduce the exact analysis. The *ArcanaPaper* class (Supplementary material), which contains methods to generate all figures in the Results section of this manuscript, is an example of this approach.

The abstraction of data and repositories in Arcana enables the same workflow implementation to be applied to datasets stored in BIDS format or XNAT repositories. A single code-base can therefore be containerized into BIDS apps or XNAT pipelines without adaptation, helping to form a bridge between the two communities of users and developers. Alternative data storage systems (Scott et al., 2011; Das et al., 2012; Book et al., 2013), can be integrated into Arcana by overriding a small number of methods from the Repository abstract base class. Repository modules could also be created for data portals such as *DataLad* (Halchenko et al., 2018) in order to take advantage of the range of platforms they support. Implementing analyses in Arcana therefore enables researchers and research groups to easily migrate their workflows between storage platforms, and not risk being locked in to a particular technology.

While Arcana was primarily developed for neuroimaging datasets, it is a general framework that could be applied to data from other fields. However, in other contexts, the subject and visit hierarchy may no longer make sense. In many cases it may be sufficient to map subjects and/or visits onto alternative concepts (e.g. for meteorological data *subjects* = *weather stations, visits = observation times*). But some cases may require a deeper data hierarchy (i.e. greater than two), which is not currently possible in Arcana.

Ensuring that the consistent versions of external tools are used throughout the analysis is important to avoid introducing biases due to algorithm updates. In systems with environment modules (Furlani, 1991) installed, Arcana can load and unload the required modules before and after each node is executed. When running Arcana within a container, environment modules and software versions can be installed inside the container giving exact control over the versions used. To these ends, a Docker container is available on Docker Hub, (hub.docker.com/r/monashbi/banana), which can be used as a base for biomedical imaging analysis containers. In future versions of Arcana, additional Environment modules could be implemented to run each pipeline node within its own container to take advantage of containers maintained by tool developers (e.g. hub.docker.com/r/vistalab/freesurfer/).

While the same tools and versions should be applied across an analysis to avoid bias, there are cases where it is desirable to rerun the same analysis with different tools substituted at various points in the workflow. In particular, when introducing new tools or upgrades to existing tools, it is important to show the effect on the final results in comparison with existing methods. Furthermore, it is typically not clear what variability between results produced by comparable tools is due to. Therefore, in the absence of *a priori* reason to favour a particular tool, perhaps the most rigorous approach is to rerun analyses with different combinations of available tools and only present results that are robust to the “analytic noise” (Maumet, 2018) they introduce. Switch parameters make it straightforward to rerun analyses in Arcana with substituted tools while controlling all other aspects of the workflow.

Arcana’s management of intermediate derivatives and provenance guarantees that the same analysis is applied across the dataset without necessarily requiring a complete rerun of the analysis. This guarantee makes it feasible to process data as it is acquired over the course of long studies, and therefore help detect any problems that might arise with the acquisition protocol when they occur. In addition, by reusing shared intermediate derivatives between analyses, such as the preprocessed DWI shared between tensor and fibre tracking workflows (Figure 11 and 12), processing time as well as time required for manual QC is minimised. Given analyses implemented in Arcana are also able to be processed on HPC clusters, they scale well to large studies.

## Conclusion

By managing the complete flow of data from/to a repository with modular components, Arcana enables complex analyses of large-scale neuroimaging studies that are portable across a wide range of research sites. The extensibility of analyses implemented in Arcana, coupled with the flexibility afforded by programmatic constructuction of pipelines, facilitates the design of comprehensive analyses by larger communities. Larger communities of developers working on the same code-base should make it feasible to capture the arcana of neuroimaging analysis in templates that can be applied to a wide range of relevant datasets.

## Supporting information

Python code for ArcanaPaper Study class and its application to a basic dataset

## Acknowledgements

The authors acknowledge the facilities and scientific and technical assistance of the National Imaging Facility, a National Collaborative Research Infrastructure Strategy (NCRIS) capability, at Monash Biomedical Imaging, Monash University. The “transparent repository” feature of Arcana was inspired by in-house software written by Parnesh Raniga while he was employed at Monash University prior to 2016.

