## Supplementary material for "A comprehensive framework to capture the arcana of neuroimaging analysis": Python code for ArcanaPaper Study class and its application to a basic dataset

#!/usr/bin/env python3
from arcana import (
    MultiStudy, MultiStudyMetaClass, SubStudySpec, ParamSpec)
from banana.study import (
    DwiStudy, T1Study, T2Study, T2starStudy, MriStudy)
from banana.plot import ImageDisplayMixin


class ArcanaPaper(MultiStudy, ImageDisplayMixin,
                  metaclass=MultiStudyMetaClass):
    """
    An Study class to perform the analyses performed in the paper
    introducing Arcana. Requires a repository containing
    T1-weighted, T2*-weighted and diffusion MRI data.
    """

    add_sub_study_specs = [
        # Sub-study to process the T1-weighted data
        SubStudySpec(
            't1',
            T1Study,
            name_map={
                't2_coreg': 't2_coreg'}),
        SubStudySpec(
            't2',
            T2Study,
            name_map={
                'coreg_ref': 't1_magnitude'}),
        # Sub-study to process the T2*-weighted data
        SubStudySpec(
            't2star',
            T2starStudy,
            name_map={'coreg_ref_brain': 't1_brain',
                      'coreg_to_atlas_mat': 't1_coreg_to_atlas_mat',
                      'coreg_to_atlas_warp': 't1_coreg_to_atlas_warp'}),
        # Sub-study to process the dMRI data
        SubStudySpec(
            'dmri',
            DwiStudy),
        # Since we are using the SWI image produced from the scanner console
        # we need to brain extract it in order to compare with QSM/vein image/
        # mask, which we do with a basic MriStudy.
        SubStudySpec(
            'swi',
            MriStudy)]

    add_param_specs = [
        ParamSpec(
            'dmri_fig_slice_offset', (6, 0, 0),
            desc=("Offset the sagittal slices of dMRI figures so they "
                  "intersect a cerebral hemisphere instead of the "
                  "midline between the hemispheres")),
        # Override default values to reduce the number of tracks in order to
        # make it easier to see the tracts in tractography figure.
        ParamSpec('dmri_num_global_tracks', int(1e5)),
        ParamSpec('dmri_global_tracks_cutoff', 0.175)]

    def vein_fig(self, save_path=None, **kwargs):
        """
        Generates an image panel containing the SWI, QSM, vein atlas,
        and vein mask

        Parameters
        ----------
        save_path : str | None
            The path to save the image to. If None the image will be
            displayed instead of saved
        """
        # Derive (when necessary) and access data SWI, QSM, vein atlas,
        # and vein mask data in the repository.
        vein_masks = self.data('t2star_vein_mask')
        qsms = self.data('t2star_qsm')
        swis = self.data('swi_brain')
        cv_image = self.data('t2star_composite_vein_image')
        # Loop through all sessions
        for swi, qsm, vein_atlas, vein_mask in zip(swis, qsms, cv_image,
                                                   vein_masks):
            # Set the saturation limits for the QSM image
            row_kwargs = [{} for _ in range(4)]
            row_kwargs[1]['vmax_percentile'] = 95
            row_kwargs[1]['vmin_percentile'] = 5
            # Display slices from the filesets in a panel using
            # method from the ImageDisplayMixin base class.
            self.display_slice_panel(
                (swi, qsm, vein_atlas, vein_mask),
                row_kwargs=row_kwargs, **kwargs)
            # Display or save to file the generated image.
            self.show(save_path, swi.subject_id, swi.visit_id)

    def fa_adc_fig(self, save_path=None, **kwargs):
        """
        Generates an image panel containing the derived FA and ADC
        images for each session in the study (typically only one)

        Parameters
        ----------
        save_path : str | None
            The path to save the image to. If None the image will be
            displayed instead of saved
        """
        # Derive FA and ADC if necessary and return a handle to the
        # data collections in the repository. Here we derive both FA and
        # ADC in a single 'data' method call so the workflows used to
        # derive them are combined.
        fas, adcs = self.data(('dmri_fa', 'dmri_adc'))
        # Loop through all sessions
        for fa, adc in zip(fas, adcs):
            # Display slices from the filesets in a panel using
            # method from the ImageDisplayMixin base class.
            self.display_slice_panel(
                (fa, adc),
                row_kwargs=({'vmax': 1.0}, {}),
                offset=self.parameter('dmri_fig_slice_offset'),
                **kwargs)
            # Display or save to file the generated image.
            self.show(save_path, fa.subject_id, fa.visit_id)

    def tractography_fig(self, save_path=None, **kwargs):
        """
        Generates dMRI tractography streamlines seeded from without the
        white matter.

        Is a bit more complex as it uses MRtrix's mrview to display and
        screenshot to file the high volume of streamlines created. Then
        reload, crop and combine the different slice orientations into
        a single panel.

        Parameters
        ----------
        save_path : str | None
            The path of the file to save the image at. If None the
            images are displayed instead
        size : Tuple(2)[int]
            The size of the combined figure to plot
        """
        # Generate and loop through tcks and FA maps for each
        # session in repository
        for tck, fa in zip(*self.data(('dmri_global_tracks',
                                       'dmri_fa'))):
            # Display generated streamlines with mrview from the
            # MRtrix package
            self.display_tcks_with_mrview(
                tcks=(tck,), backgrounds=(fa,),
                offset=self.parameter('dmri_fig_slice_offset'),
                **kwargs)
            # Display or save to file the generated image.
            self.show(save_path, tck.subject_id, tck.visit_id)


if __name__ == '__main__':
    import os
    import os.path as op
    import logging
    from argparse import ArgumentParser
    from arcana import (
        BasicRepo, SingleProc, StaticEnv, FilesetInput)
    from arcana.utils import parse_value
    from banana.file_format import dicom_format, zip_format

    logger = logging.getLogger('arcana')
    logger.setLevel(logging.INFO)
    handler = logging.StreamHandler()
    formatter = logging.Formatter("%(levelname)s - %(message)s")
    handler.setFormatter(formatter)
    logger.addHandler(handler)

    # Set up parser to parse arguments passed to the script
    parser = ArgumentParser(
        "A script to produce figures for the Arcana manuscript")
    parser.add_argument('data_dir',
                        help="The directory containing repository")
    parser.add_argument('--work_dir', help="The work directory",
                        default=op.join(op.expanduser('~'),
                                        'arcana-paper-work'))
    parser.add_argument('--fig_dir',
                        help="The directory to store the figures",
                        default=op.join(op.expanduser('~'),
                                        'arcana-paper-fig'))
    parser.add_argument('--t1', default='.*mprage.*',
                        help="Pattern to match T1-weighted scan")
    parser.add_argument('--t2', default='.*t2.*',
                        help="Pattern to match T1-weighted scan")
    parser.add_argument('--t2star_chann', default='.*channels.*',
                        help=("Pattern to match separate channels from the "
                              "head coil for the T2*-weighted scan in "
                              "separate NiFTI files within a single "
                              "directory"))
    parser.add_argument('--swi', default='.*(swi|SWI).*',
                        help="Pattern to match SWI scan in DICOM format")
    parser.add_argument('--dwi', default='.*diff.*',
                        help="Pattern to match dMRI scan")
    parser.add_argument('--distort', default='.*distortion_correction.*',
                        help="Pattern to match dMRI reverse PE ref. scan")
    parser.add_argument('--parameter', '-p', nargs=2, action='append',
                        metavar=('NAME', 'VALUE'), default=[],
                        help="Parameters to pass to the study")
    parser.add_argument('--reprocess', action='store_true', default=False,
                        help=("Reprocess derivatives on mismatches between "
                              "requested parameters and stored provenance"))
    args = parser.parse_args()

    # Make figure directory
    os.makedirs(args.fig_dir, exist_ok=True)

    # Instantiate the ArcanaPaper class in order to apply it to a
    # specific dataset
    paper = ArcanaPaper(
        # Name for this analysis instance
        'arcana_paper1',
        # Repository is a simple directory on the local file system
        repository=BasicRepo(args.data_dir),
        # Use a single process on the local system to run pipelines
        processor=SingleProc(args.work_dir, reprocess=args.reprocess),
        # Use the software installed and on the curren PATH (i.e. no Modules)
        environment=StaticEnv(),
        # Match names in the data specification to filenames used
        # in the repository
        inputs={
            't1_magnitude': FilesetInput(pattern=args.t1, dicom_format),
            't2_magnitude': FilesetInput(pattern=args.t2, dicom_format),
            't2star_channels': FilesetInput(pattern=args.t2star_chann,
                                            zip_format),
            't2star_header_image': FilesetInput(pattern=args.swi,
                                                dicom_format),
            't2star_swi': FilesetInput(pattern=args.swi, dicom_format),
            'swi_magnitude': FilesetInput(pattern=args.swi, dicom_format),
            'dmri_magnitude': FilesetInput(pattern=args.dwi, dicom_format),
            'dmri_reverse_phase': FilesetInput(pattern=args.distort,
                                               dicom_format)},
        # Set parameters of the study
        parameters={n: parse_value(v.strip()) for n, v in args.parameter})

    # Derive required data and display them in a single step for each
    # figure.
    paper.vein_fig(op.join(args.fig_dir, 'veins.png'))
    paper.fa_adc_fig(op.join(args.fig_dir, 'fa_adc.png'))
    paper.tractography_fig(op.join(args.fig_dir, 'tractography.png'))
